# Investigating Honey Bee Pollen Foraging Patterns Across a Season Using DNA Metabarcoding

**DOI:** 10.1101/2024.10.11.617889

**Authors:** Sydney N. Larsen, Joshua G. Steffen, Heather M. Briggs

## Abstract

The western honey bee, or *Apis mellifera*, is a vital pollinator, essential for both ecosystems and agriculture worldwide. As human influences on these bees become more pronounced, understanding their pollen foraging behavior is increasingly important. Pollen provides crucial nutrients for *A. mellifera* such as protein and lipids that are necessary for hive health and prosperity. Pollen DNA metabarcoding allows for the longitudinal analysis of the floral sources of corbicular (bee-collected) pollen. In this study we employ the use of metabarcoding and Oxford Nanopore Sequencing technology to analyze the floral composition of pollen samples collected on a weekly basis from two closely related hives. In total, we identified 74 distinct plant taxa (at the genus level) between the two hives. Although both hives had equivalent values of taxonomic richness and diversity, the majority of taxa identified were unique to the individual hives. This indicates a large degree of variation in pollen foraging despite the hives’ relative proximity. In addition to this interhive analysis, we also analyzed changes in pollen origin of a single hive across the course of five months. We identified 59 distinct plant taxa within samples from this hive whose presence and relative abundance differ drastically on a weekly basis with no taxa being present in all samples across the observed time period. In addition, even though there was a median of 8 to 9 taxa detected in each sample, each sample was composed primarily of 1 to 2 taxa indicating a degree of foraging preference that changes on a regular basis. Further quantitative and qualitative comparisons completed in this study demonstrate the degree of variation in pollen foraging patterns both throughout the season and between hives.

## Introduction

*Apis mellifera*, commonly known as the western honey bee, is one of the most predominant bee pollinator species in the world. *A. mellifera* were observed in 88% of pollination networks within their native ranges, contributed to an average of 13% of total floral visitation, and were the predominant floral visiter (≥50%) to 17% of plant taxa even in pollination systems outside of their local range (Hung et al., 2018). Agriculture is also highly dependent on this species with 80% of crop pollination being attributed to *A. mellifera* due to their ideal body size, high attentiveness, and ability to boost the fitness of local floral populations (Idrees et al., 2023). The significance of this species to global agriculture and ecosystems highlights the need to understand the delicate relationships between these insects and their environment.

*A. mellifera* pollination behavior is a secondary consequence of their need for nutrient and energy acquisition. They visit flowers in order to obtain crucial nutrients from nectar and pollen which they transport back to the colony. During a single foraging trip, honey bees can visit up to five hundred flowers to obtain a sufficient amount of resources (Winston, 1995). Nectar serves as the major source of energy for the colony, while pollen is the bees’ source of protein, lipids, sterols, vitamins, and minerals and plays a crucial role in larvae development (Dharampal et al., 2019; Thornburg, 1970). Despite flowering plants being sources of pollen and nectar simultaneously, the majority of workers specialize in predominantly pollen or nectar collection. Some studies suggest that pollen collection is largely completed by more experienced foragers (Klein et al., 2019; Younas et al., 2022).

Pollen foragers aggregate individual pollen grains from across their bodies into corbiculae on their legs using regurgitated nectar (Prado et al., 2023). Upon returning to the hive, the collected pollen is packed into brood comb by younger worker bees and chemically altered by the addition of nectar, honey, and glandular secretions to form the stored form of pollen called bee bread, the primary energetic nutritional resource for the colony (Herbert & Shimanuki, 1978).

The quality of pollen resources can dictate the foraging behavior of *A. mellifera*. Foraging bees have been observed selecting pollen based on specific hive needs that correlate with different nutritional values, such as lipid and amino acid composition of these resources rather than per quantity (Cook et al., 2003; Day et al., 1990; Hendriksma & Shafir, 2016; Zarchin et al., 2017). This dedication to pollen quality explains why *A. mellifera* pollen foragers have been observed to have preferences for pollen that are distinct and more selective than corresponding nectar preferences (Do, Y., et al., 2022). Furthermore, honey bees tend to forage on fewer species of plants to obtain pollen resources at one time relative to nectar, but the species of which they forage on vary across both hives and time (Coffey & Breen, 1997; McMinn-Sauder et al., 2022). While honey (the stored form of nectar) is stored over long periods of time, bee bread is consumed relatively quickly, 95% of all collected pollen is consumed within two weeks (Roessink & van der Steen, 2021). This rapid consumption of pollen means that, at any given time during the foraging season, an absence of high-quality pollen could have detrimental impacts on the development, productivity, and reproduction of the colony (PaдB et al., 2014).

Research examining the origin of corbicular pollen has classically relied on microscopic identification which is time-intensive, expensive, conditionally-biased, and generally results in shallow pollen identification (Prudnikow et al., 2023). An alternative to microscopic identification is DNA metabarcoding which allows for the identification of pollen using relatively short, but taxonomically distinct DNA sequences. This technology coupled with next generation sequencing (NGS) allows for the sequencing of organisms present in high-complexity DNA samples. DNA metabarcoding approaches facilitate the study of the foraging behavior of multiple hives from pollen samples by increasing the number of taxa identified and decreasing the time and specialized training required using microscopy (Keller et al., 2015; Smart et al., 2017).

Many recent examples of using DNA metabarcoding as a tool to study foraging utilize bee bread rather than corbicular pollen to understand the dietary impacts of nutrition on the hive largely because bee bread is more bioactive and has higher percentages of proteins, lipids, fibers, etc due to the lactic acid fermentation process (Aylanc et al., 2021; Lu et al., 2021; Tamizi et al., 2019). In addition to the increased bioactivity of bee bread, this form of pollen also differs in its plant composition due to the presence of bee saliva and nectar (Bakour et al., 2022). While analyses of bee bread provide insights into pollen collection, the presence of nectar may prevent accurate descriptions of pollen foraging behavior.

Here we employ the use of the DNA metabarcoding approach to corbicular pollen to obtain a description of pollen foraging behavior throughout a hive’s foraging season. Our work also examines variation in foraging behavior by comparing two adjacent hives. The use of corbicular pollen instead of bee bread fosters the analysis of pollen foraging behavior while limiting the impact of cross-contamination from nectar. The information gained from this analysis will serve as a baseline of season-wide foraging trends in diversity and plant taxa visited and elevate our understanding of the role that hive identity plays in pollen foraging.

## Methods

### Sampling Locations

This investigation employs the use of two distinct hives located on the University of Utah campus in Salt Lake City, UT (**Figure 1)**. These hives are maintained by the University of Utah Beekeepers Association which provided access during the 2023 foraging season.

**Figure 1:**
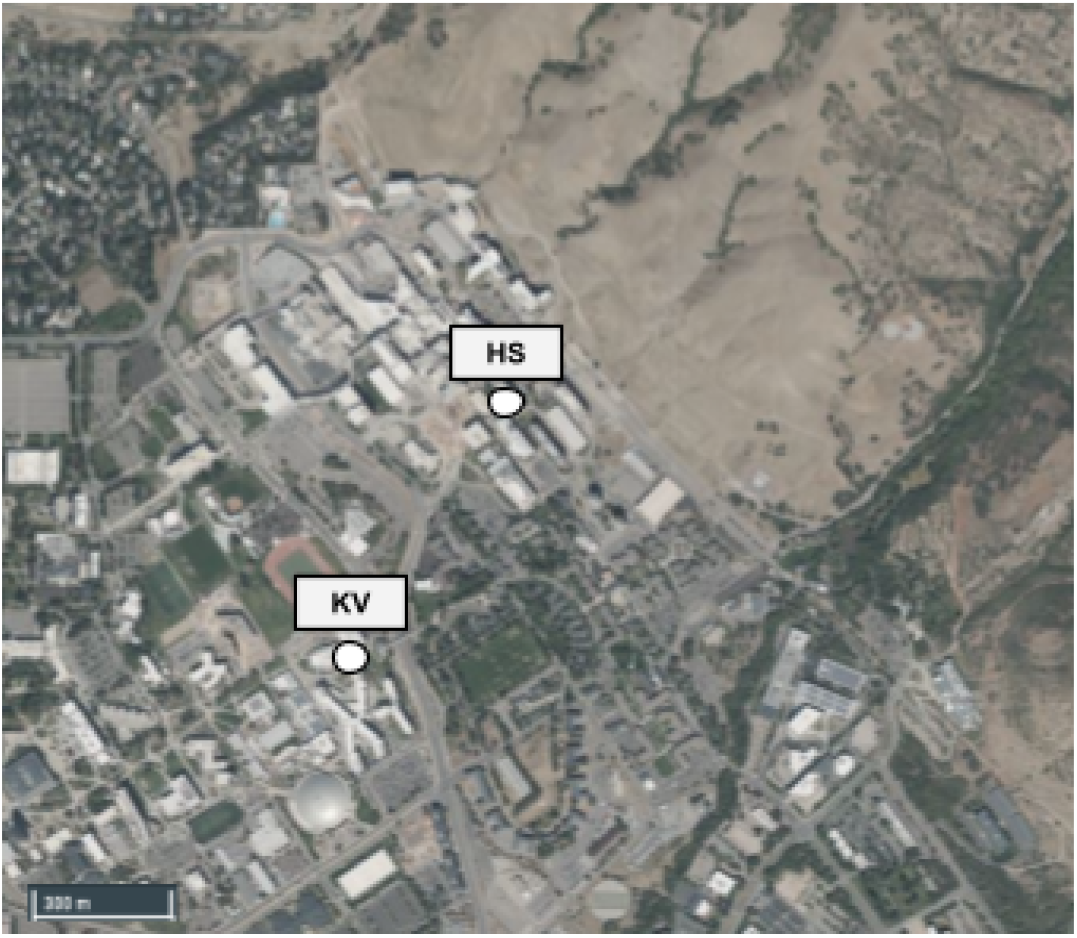
Location of the Two Hives. University of Utah campus with the two locations of the hives indicated with white dots and acronyms. The full names and coordinates of each hive are found above.

Hive one is located within a plant-rich, landscaped area directly adjacent to the Health Science Education Building (40.76952° N, 111.83413° W) and is here on out referred to as the HS hive. While the walkway next to this building is highly traveled, since the hives are tucked within the vegetation, there is little direct human interaction between humans and the hives and bees themselves. This hive is located along the foothills of the Wasatch Mountain Range and therefore is exposed to a variety of native flora and fauna.

Hive two is located next to a high-trafficked walkway and road on the corner of the Kahlert Village Dorms (40.76465° N, 111.83789° W) and is referred to as the KV hive hereafter. The KV hive itself is located in the shade of a large tree. While there is a low amount of vegetation immediately surrounding the hive, ornamental flora and fauna are located only yards away. Hive two is found on the north side of the Health Science Education Building and is henceforth referred to as HS (40.76952° N, 111.83413° W).

The two hives surveyed in this study are separated by 0.63 km. The mean foraging distance for Apis mellifera is approximately 5.5 km with over 50% of bees foraging at a distance greater than 6 km (Beekman & Ratnieks, 2000). This large foraging range means that although the two hives are immediately surrounded by unique landscapes, they are well within a comparable foraging area.

### Pollen Collection

Corbicular pollen was collected by employing the use of pollen traps (HunterBee, 10 Frame Removable Pollen Trap). These traps, when set, lower a grid that covers the front entrance of the hive. This forces the bees, upon returning from a foraging trip, to squeeze through these small openings scraping the pollen pellets from the bees’ corbicula into a basket that is set underneath the trap. This collection basket is covered in sanitized (70% ethanol) aluminum foil. The addition of freshly sanitized foil prior to every collection prevented contamination from week to week.

Pollen collection occurred weekly, beginning on June 6, 2023 and ending on October 17, 2023. The daily timing of pollen collection was decided in accordance with findings from an article from the Institute of Apicultural Research, Chinese Academy of Agricultural Sciences published in 2019 which indicates the proportion of viable pollen carried by A. mellifera increases throughout the morning from 9:30 am to 2:00 pm then decreases steadily in the afternoon (Delaplane et al., 2013).

For each collection day, metrics regarding the quantity of pollen collected, timing of pollen collection, and weather conditions were recorded **(Table 1)**. Once the pollen was collected, it was then weighed, imaged, and stored in individual 50 mL sterile polypropylene conical centrifuge tubes at -80°C.

**Table 1:** Pollen Collection Description. The measured weather conditions comprises the daily average: temperature, humidity, precipitation, and air pressure, and air quality index. The temperature, humidity, precipitation, and air pressure values were obtained from the Weather Underground, Salt Lake City International Airport Station history records; the air quality index was obtained from IQair.com Salt Lake City Station.

| Collection | Weather Conditions |  |  |  |  | Pollen Information |  |  |  |  |
| --- | --- | --- | --- | --- | --- | --- | --- | --- | --- | --- |
| Date | Temp. (°C) | Humidity (%) | Precip. (in) | Pressure (in) | Air Quality (US AQI) | HIVE | Quantity of Pollen (g) | Trap Set | Collected | Duration |
| 06/06/23 | 23.6 | 40.8 | 0.00 | 25.6 | 16 | HS | 4.46 | 11:17 | 14:07 | 2:50 |
|  |  |  |  |  |  | KV | 1.69 | 10:59 | 13:40 | 2:41 |
| 06/13/23 | 20.6 | 45.1 | 0.00 | 25.7 | 8 | HS | 0.71 | 9:29 | 11:58 | 2:29 |
|  |  |  |  |  |  | KV | 0.21 | 9:54 | 12:21 | 2:27 |
| 06/20/2023 | 16.6 | 56.6 | 0.03 | 25.7 | 24 | HS | 0.77 | 10:18 | 13:29 | 3:11 |
|  |  |  |  |  |  | KV | 1.45 | 10:24 | 13:48 | 3:24 |
| 06/27/2023 | 23.9 | 36.7 | 0.00 | 25.6 | 16 | HS | 14.58 | 10:27 | 13:23 | 2:56 |
|  |  |  |  |  |  | KV | 6.76 | 10:45 | 13:45 | 3:00 |
| 07/06/2023 | 27.2 | 25.9 | 0.00 | 25.6 | 21 | HS | 0.34 | 10:25 | 13:40 | 3:15 |
|  |  |  |  |  |  | KV | 3.75 | 10:40 | 14:00 | 3:20 |
| 07/11/2023 | 30.4 | 14 | 0.00 | 25.7 | 33 | HS | 10.59 | 10:25 | 13:40 | 3:15 |
|  |  |  |  |  |  | KV | 0.19 | 10:40 | 14:00 | 3:20 |
| 07/18/2023 | 30.3 | 29.1 | 0.00 | 25.8 | 35 | HS | 0.18 | 10:15 | 13:15 | 3:00 |
|  |  |  |  |  |  | KV | 2.27 | 10:23 | 13:30 | 3:07 |
| 07/25/2023 | 29.0 | 41.6 | 0.03 | 25.8 | 51 | HS | 35.49 | 10:13 | 13:30 | 3:17 |
|  |  |  |  |  |  | KV | 3.98 | 10:32 | 13:49 | 3:17 |
| 08/01/2023 | 27.7 | 39.3 | 0.00 | 25.8 | 39 | HS | 0.1 | 10:06 | 13:06 | 3:00 |
|  |  |  |  |  |  | KV | 0.42 | 10:30 | 13:20 | 2:50 |
| 08/08/2023 | 22.9 | 45.2 | 0.01 | 25.7 | 47 | HS | 1.51 | 10:16 | 13:32 | 3:16 |
|  |  |  |  |  |  | KV | 0.22 | 10:34 | 13:54 | 3:20 |
| 08/15/2023 | 29.4 | 33.5 | 0.00 | 25.8 | 45 | HS | 9.95 | 10:12 | 13:31 | 3:19 |
|  |  |  |  |  |  | KV | 0.21 | 10:34 | 13:53 | 3:19 |
| 08/22/2023 | 21.2 | 69.3 | 0.00 | 25.8 | 29 | HS | 1.58 | 10:08 | 13:42 | 3:34 |
|  |  |  |  |  |  | KV | 0.36 | 10:19 | 13:58 | 3:39 |
| 08/29/2023 | 29.3 | 26.5 | 0.00 | 25.7 | 39 | HS | 15.62 | 10:15 | 13:07 | 2:52 |
|  |  |  |  |  |  | KV | 0.33 | 10:32 | 13:33 | 3:01 |
| 09/12/2023 | 23.6 | 35.2 | 0.00 | 25.7 | 20 | HS | 25.16 | 10:02 | 13:01 | 2:59 |
|  |  |  |  |  |  | KV | 2.42 | 10:10 | 13:11 | 3:01 |
| 09/19/2023 | 22.2 | 38.5 | 0.00 | 25.7 | 22 | HS | 12.71 | 9:55 | 12:57 | 3:02 |
|  |  |  |  |  |  | KV | 0.32 | 10:10 | 13:19 | 3:09 |
| 09/26/2023 | 22.7 | 28.6 | 0.00 | 25.8 | 26 | HS | 25.96 | 9:45 | 12:57 | 3:12 |
|  |  |  |  |  |  | KV | 0.34 | 10:00 | 13:16 | 3:16 |
| 10/03/2023 | 11.4 | 84 | 0.39 | 25.8 | 24 | HS | 0 | 9:54 | 12:47 | 2:53 |
|  |  |  |  |  |  | KV | 0 | 10:10 | 13:01 | 2:51 |
| 10/17/2023 | 13.4 | 68.1 | 0.00 | 25.8 | 38 | HS | 4.29 | 9:51 | 13:08 | 3:17 |
|  |  |  |  |  |  | KV | 0.16 | 10:10 | 13:22 | 3:12 |

### DNA Extraction

In order to extract DNA from the pollen samples, one gram of pollen (0.7g for low-quantity samples) was solubilized in 400µl of sterile, distilled water. From this diluted pollen solution, 200mg of pollen (400µL of the diluted solution) was then used as the input for DNA extraction. Extractions were completed using the Qiagen DNeasy® Plant Pro Kit (Qiagen, ID: 69206) with slight modifications. These modifications include adding 650µl of APP Buffer as opposed to 500µl and adding 50µl of EB Buffer which is the minimum amount recommended.

### Metabarcoding and Multiplexing PCR

After extracting the DNA, polymerase chain reactions (PCR) were used to amplify the chloroplast trnL (UAA) gene via the trnLC/trnLD primer pair. To aid with demultiplexing in future steps, one of eight unique indexes were attached to each primer, four associated with forward primers and four associated with reverse primers **(Table S1)**.

The PCR reactions were accomplished in accordance with Phusion High Fidelity 2x Master Mix (Thermo Fisher Scientific) guidelines. This protocol consisted of: 15µL of Phusion High Fidelity 2x Mastermix, 3 µL of forward primer, 3µL of reverse primer, 2 µL of 25 ng/µL DNA template stock, and 7 µL of nuclease free water for a final PCR volume of 30 µL.

After testing a range of PCR cycling protocols, the finalized cycling conditions consisted of: one cycle of 98°C for 30s, 22 cycles (minimum number of cycles for sufficient amplification) of 98°C for 10s, 50°C for 30s, and 72°C for 30s. This cycling is followed by a final extension at 72°C for 10 minutes. PCR products were then tested on a 1% agarose gel.

### Library Preparation and Oxford Nanopore Technologies Sequencing

In order to prepare the PCR samples for sequencing, we used the Qiagen QIAquick PCR Purification using a Microcentrifuge protocol (Qiagen, 28106). After completing the purification, the concentration of DNA was recorded using the Qubit^™^ Fluorometer 4.0 and the Qubit^™^ dsDNA HS protocol (Thermo Fisher Scientific, Q32854).

During one sequencing run, sixteen PCR samples (each containing a unique combination of forward and reverse indexed primers) were multiplexed at equimolar ratios to reach a final value of 100 fmols. The multiplexed DNA library were then prepared in accordance with ONT protocols for the SQK–LSK114 sequencing kit, adapted for the use of MinION Flongle Flow Cell (R10.4) (Oxford Nanopore Technologies, SQK-LSK114 & New England Biolabs, NEB #E7672) with slight alterations to incubation time (30 min at 20°C and 30 min at 65°C instead of 5 mins intervals) and adding the AMPure XP beads at the same volume as the amplicon (50 µl) rather than the recommended 20 µl. Following preparation, 9-10 fmols of DNA were sequenced using the MinION (Mk1C) sequencing device over a 36 hour period. The reads were exported as POD5 files.

### Demultiplexing and Taxa Identification

The sequences present in POD5 files were basecalled using the high-performance, open source basecaller, dorado (v 0.6.0) (https://github.com/nanoporetech/dorado/tree/release-v0.6.0) and filtered to only contain sequences with a minimum read length of 200 bp and a minimum Quality Score (Q-score) of 8. The sequences were then demultiplexed via their unique barcode combination using minibar.py (v 0.24, written in Python 2.7) while simultaneously removing all barcodes and indexes at a primer editing distance of e9. The subsequent demultiplexed FASTQ files were converted into FASTA files which were then compared against a custom trnL-c/trnL-d database utilizing the NCBI Chloroplast database (downloaded on May 12th, 2024) and scripts provided by R. Green (https://github.com/robertgreenhalgh/stand). We first searched terms within the entire NCBI Blast database, “ biomol_genomic[PROP] AND is_nuccore[filter] AND chloroplast[filter]”, to compile a collection of only chloroplast sequences. This database of chloroplast sequences was further reduced to a collection of trnL sequences which contained only GGGCAATCCTGAGCCAA (trnL-forward) and CCATTGAGTCTCTGCACCTATC (trnL-reverse) sequences. This final database contained 18,462 sequences that are Viridiplantae *trnL* specific with a maximum of three total primer mismatches (maximum of two mismatches per primer) and amplicon sequence length between 8 and 175 bp.

Taxonomic units are presented at the genus level due to the lack of species level specificity of the trnL primer (Taberlet et al., 2007). To remove any possible misaligned reads due to contamination, sequencing error, or taxonomic identification error, taxa with less than 0.05% of the sample’s total read number and/or less than 10 total reads were omitted. In addition, any sample with less than 2500 total reads after performing this filtering were omitted to prevent shallow sequencing from impacting the results as seen in Leponiemi et al., 2023.

### Statistical Analysis

Statistical analyses and all corresponding figures creation were generated using R(V 4.4.0) with functions present in the vegan package (V 2.6-4) (Oksanen, J., et al., 2013).

#### Menhinick’s Richness Index

In order to account for the correlation between the number of sequences and number of taxa identified, the Menhinick’s Index (D) represents taxa richness as the number of distinct species in a sample (n) divided by the square root of the total number of individuals within the sample (N).

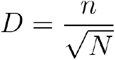

#### Shannon Diversity Index

The Shannon Diversity Index considers both taxa richness and evenness and is based on the uncertainty of the identity of a species chosen at random. A larger diversity within a sample results in high uncertainty regarding the identity of a random taxa (larger H’ value). The Shannon Diversity Index can be represented as

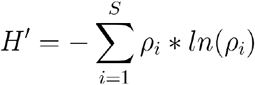

Where ρ_i_ = proportion of individuals of species i (n_i_ / N) and S is the taxa richness. The Shannon Diversity of each sample is calculated using the “ diversity” function from the vegan package (“ shannon” is the default analysis method for this function).

The following analyses were performed using Hellinger Transformed Data which was created from the RRA data using the function “ decostand” from the vegan package using the method “ hellinger”.

data_hellinger <-decostand(counts_RRA, method = “ hellinger”)

#### Nonmetric Multidimensional Scaling

In order to determine how different temporal seasons and hives differ and group based on plant composition, we performed nonmetric multidimensional scaling (NMDS) using the vegan package function “ metaMDS” in 2-dimensions (k = 2) on the Hellinger-transformed data.

The distance matrix was created using the function “ vegdist” from vegan package and the Hellinger transformed data (using the ‘euclidean’ method). In order to test if these groups are statistically distinct, the distance of the group members to the group centroid are subjected to the “ anova” function (vegan package)

hellinger_dist <-vegdist(data_hellinger, method = ‘euclidean’)

For the following analyses, a P-value less than 0.05 will be considered statistically significant for the purposes of this study.

#### PERMDISP

PERMDISP is the multivariate counterpart of the Leven homogeneity test for multivariate and tests the null hypothesis that there is no difference in the distribution between groups. In order to perform an analysis of multivariate homogeneity of group diversions the following code

permadisp_results <-betadisper(hellinger_dist, df_RRA$season)

#### PERMANOVA

Permutational multivariate ANOVA (PERMANOVA) is a non-parametric alternative to a multivariate ANOVA test and is described as a geometric partitioning of multivariate variation (Anderson, 2017). This semiparametric method of performing an ANOVA test allowed us to formally partition our multivariate, non-normal data. We performed a PERMANOVA using the vegan package following this function structure

permanova_results <-adonis2(data_hellinger ∼ df_RRA$season)

#### PAIRWISE PERMANOVA

Pairwise PERMANOVAs were performed on Hellinger Transformed distance data to partition the multivariate data between the each of the three temporal subgroups of the HS hive samples with bonferroni adjustment (https://gist.github.com/mcgoodman).

## Results

### Sequencing Results

Pollen was collected from two hives on the University of Utah campus, the HS hive, located near a non-developed area of the Wasatch foothills, and the KV hive situated closer to the center of campus. From the eighteen total collection days, thirty-four samples of pollen were collected with the weight of pollen ranging from 0.01g to 25.96g. Out of these thirty-four samples, twenty-one (HS hive, n=14 & KV hive, n = 7) met the set quality standards for the sample (weight ≥ 0.7g and total number of reads ≥ 2,500) (see **Table 1** for weather conditions and pollen quantity details). Among these twenty-one samples, there were a total of 265,368 sequencing reads obtained with 261,214 of those reads (HS hive, 143,055 reads & KV hive, 118,159 reads) meeting the set parameters for consideration of the individual taxonomic units (relative read abundance (RRA) ≥ 1% & n ≥ 10 per sample). The number of reads per individual sample ranged from 2,541 to 46,217 reads, with a median of 4,293 reads.

### Seasonal Variation

The HS hive was sampled on a weekly basis to determine the variation in foraging behavior throughout the duration of a single foraging season. Due to variations in the quantity of pollen collected by the hive, six of the eighteen weeks resulted in insufficient pollen quantity (<0.7g).

Throughout the fourteen HS pollen samples, a total of 59 distinct taxonomic units were identified at the genus level **(Table S2)**. A median of 8 taxa were detected per sample with the minimum number of taxa being 4 (HS 10/17) and the maximum being 18 (HS 6/20).

The majority of the taxa identified, 34 of the 59 taxa, had a maximum relative abundance of less than 5%. No taxa were identified in all fourteen samples with the most frequently sequenced taxa being *Taraxacum* which was present in 10 of the 14 samples. *Taraxacum* also had the second largest maximum RRA (79.82% on 10/17) behind Agastache which composed 80.23% of the 07/25 sample. Other prominent plant genera identified were *Spyridium (max. RRA, 78.98%), Chrysanthemum (max. RRA, 70.08%), Erigeron (max. RRA, 52.04%), Ulmus (max. RRA, 45.83%), and Gleditsia (max. RRA, 45.31%)*. The composition of each sample based on RRA values is shown in **Figure 2a-b**.

**Figure 2:**
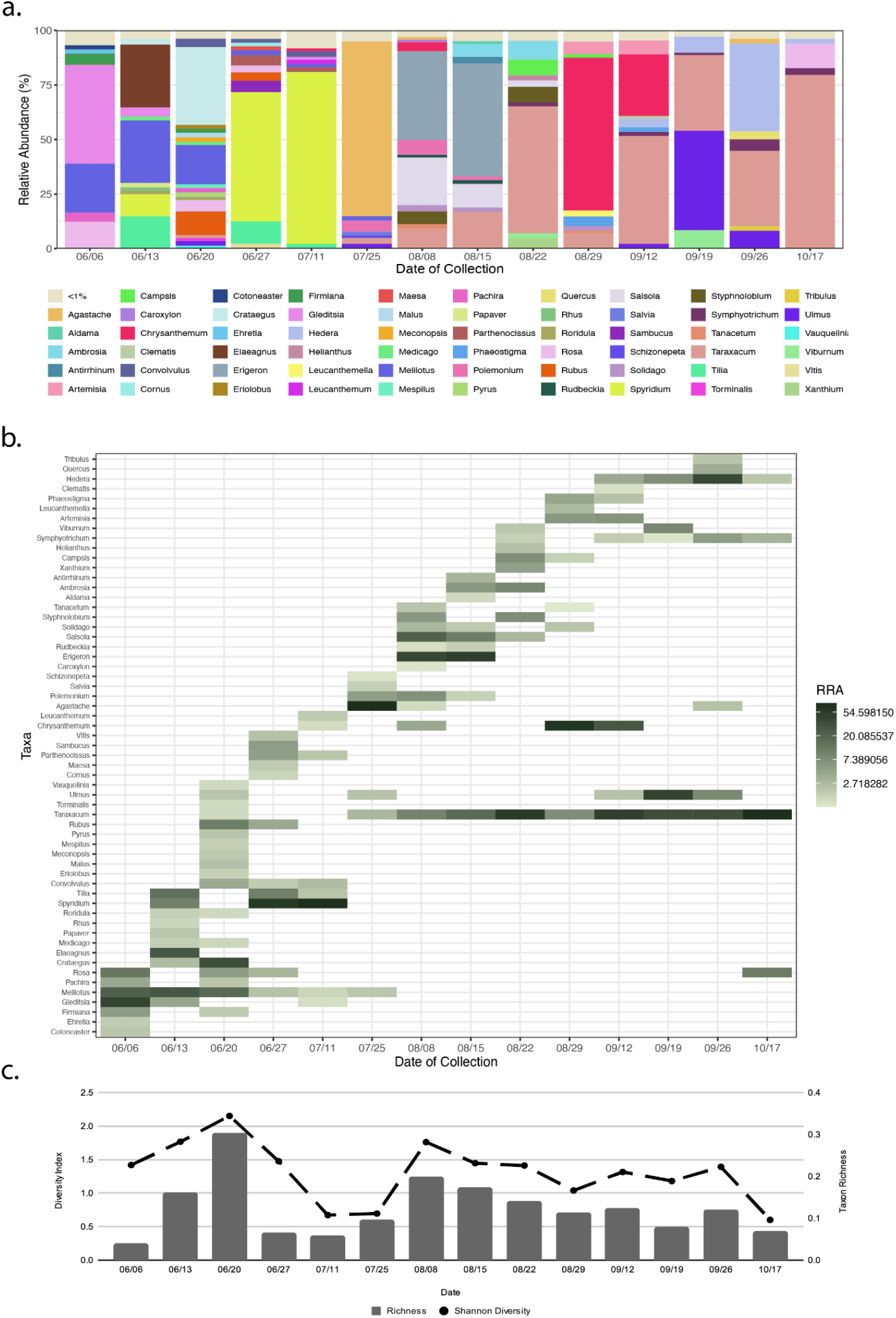
The HS Hive’s Pollen Composition over the Foraging Season. (a) Stacked bar charts showing the taxon composition (at the genus level) of plants identified in fourteen corbicula pollen samples collected from the HS hive. Taxa accounting for < 1% of individual sample’s reads are grouped as “ <1%”. (b) Heatmap expressing the RRAs of taxa on each collection day. (c) Diversity indexes of HS samples throughout the duration of the foraging season. The Shannon-Wiener Index is indicated by the dashed line (left axis), and the Menhinick’s Richness Index is indicated by the column (right axis).

The variation in the composition of pollen samples throughout the season revealed similar patterns in key diversity metrics. The taxa richness varied throughout the season with local minimums occurring on 7/11 (0.055) and 10/17 (0.066) and local maximums on 6/20 (0.30) and 8/08 (0.195). The Shannon diversity index followed a similar pattern with relatively high levels of diversity on 6/20 and 8/08 (2.15 & 1.76, respectively) and a large dip in diversity on 07/11 (0.67). Following peaks on 08/08, both diversity indexes followed a trend of steady decline for the remainder of the season with Shannon diversity hitting its absolute minimum on the last collection day of the season (10/17, 0.60) **(Figure 2c)**.

In order to analyze long-term changes in corbicular pollen composition across the foraging season, the samples were grouped into three temporal categories each representing a five-to six-week time period throughout the collection timeline: Early, Mid, and Late seasons. Samples collected from June 6th to July 11th (n = 5, spanning a 5-week period) were grouped as “ Early Season” ; samples from July 25th to August 29th (n = 5, spanning a 5-week period) are grouped into “ Mid Season”, and samples from September 12th to the end of the foraging season (October 17th) (n = 4, spanning a 6-week period) are considered “ Late Season”. This seasonal grouping closely aligns with hierarchical clustering done on this samples which shows that the Early Season samples form a defined seasonal bin while the Mid and Late Seasons have a higher degree of similarity **(Figure S1)**

The Early Season has the highest number of distinct plant taxa identified (33) with the late season containing the lowest number of taxa (13). Despite the disparity between the number of taxa identified in each sub-season, the plant composition of the three temporal seasons does not differ significantly in their dispersion (PERMADISP, Pr(>F) = 0.1302 and F-value = 2.4683).

Only 3 taxa (*Chrysanthemum, Taraxacum*, & *Ulmus*) were identified within all three seasons. There were 7 taxa present in two of the three seasons: 1 taxon (*Melilotus*) in the early and mid seasons, 1 taxon (*Rosa*) in the early and late seasons, and 5 taxa (*Agastache, Artemisia, Phaeostigma, Symphyotrichum*, & *Viburnum*) present in both mid and late seasons **(Figure 3a)**. Despite this partial overlap in plant taxa between the three temporal seasons, the relative abundance of these shared taxa differs dramatically from season to season **(Figure 3b)**. For instance, *Taraxacum*, which was present in all three seasons, is present in only one Early Season sample (06/20) and encompasses only 1.47% of that sample (0.28% of the total Early Season). Alternatively, *Taraxacum* is present in all nine Mid and Late Season samples and makes up 18.9% and 49.7% of the two seasons’ total reads, respectively. This taxonomic abundance difference as well as others explains the significant difference observed amongst the three seasons’ plant composition (PERMANOVA, Pr(>F) = 0.001 and F = 4.415) (Figure CCC). The early season differs significantly from both the mid and late seasons (Pairwise PERMANOVA w/ Bonferroni Adjustment, p = 0.036 and p = 0.024, respectively) while the mid and late seasons do not differ significantly (p = 0.138) from each other in their composition **(Figure 3c)**.

**Figure 3:**
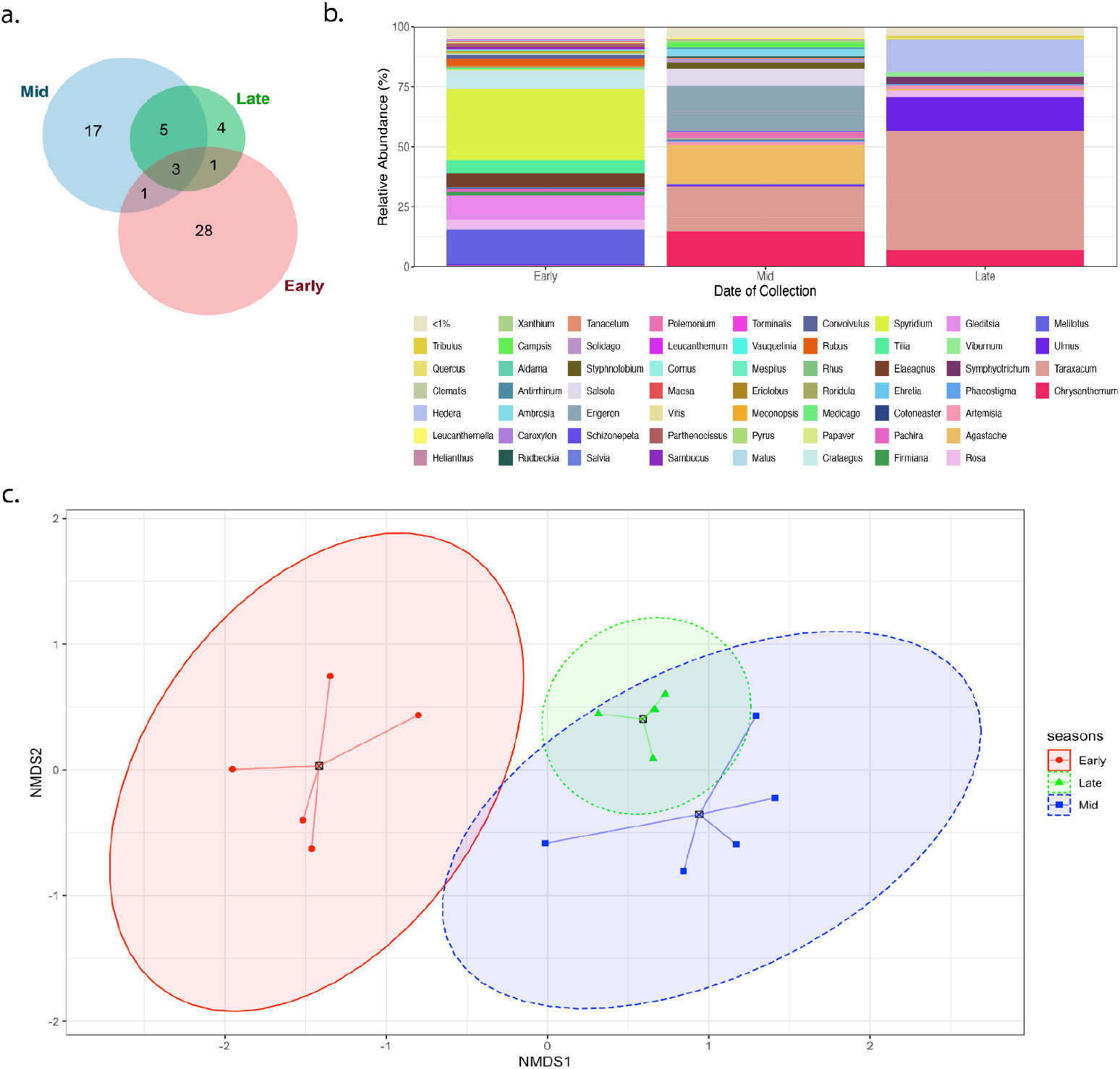
The Analysis of HS Hive’s Samples Divided into Three Temporal Seasons. (a) Number of shared and unique plant genera identified in Early Summer (red, n = 4), Mid Summer (blue, n = 5), and Late Summer (green, n = 5). (b) Stacked bar charts showing the taxon composition (at the genus level) from pooled samples for each of the three seasons. The RRA of each taxon is determined by the pooling of all samples from their respective seasons (Early = 6/6 to 6/27, Mid = 7/11 to 8/22, and Mid = 8/29 to 10/17). Taxa accounting for <1% of reads in each hive are grouped as “ <1%”. (c) This non-metric multidimensional scaling (NMDS) plot is based on Hellinger Dissimilarity Index Data and ellipses show the 95% confidence level for the three seasons.

#### Interhive Variation

In addition to sampling from the HS hives throughout the season, we collected seven samples from another geographically close hive (KV). The date of collection for these samples ranged from June 6th to September 12th. This hive is located 0.63 km away from the HS hive, well within *A. mellifera*’s mean foraging distance of 5.5 km (Beekman & Ratnieks, 2000). Throughout these seven samples, a total of 33 plant taxonomic units (at the genus level) were identified **(Table S3)**. This hive had a median of 9 distinct plant taxa per sample (ranging from 4 taxa on 09/12 to 13 taxa on 06/27). Of the 33 total genera identified, 22 genera identified throughout the KV samples had maximum RRA’s less than 10% leaving only 11 taxa to make up the majority of the KV hive samples’ composition. Similarly to the HS hive, no plant genus was found in all 7 samples. *Parthenocissus* was the most frequently observed (5 of the 7 samples) but only had the fourth highest maximum RRA (60.35%). Along with *Parthenocissus, Solidago* (max. RRA, 87.61%), *Ulmus* (max. RRA, 82.45%), and *Tilia* (max. RRA, 68.94%) were the only taxa’s that had maximum relative read abundance values over 50% while each being identified in only one to three of the seven total KV samples.

We then compared the composition of all KV samples to all HS samples in order to determine the difference in foraging behavior between these two hives. Among the two hives, 74 distinct taxa were identified, and only eighteen taxa were present in both hives. Forty-one taxa were found exclusively in the HS hive samples, and fifteen were unique to the KV hive **(Table S4)**.

Predominant genera at each sampling time varied significantly between hives. For instance, *Spyridium* (the most abundant plant genera found within the HS samples) composes 26.14% of the total HS taxa identified but was not identified within the KV samples (above the >1% cutoff). The 18 shared taxas’ RRAs also range based on the hive. For instance, *Tilia* (the most abundant taxa in KV hive) composes 21.32% of KV reads but only made up 3.07% of the HS hive’s total plant composition **(Figure 4a)**. The partial overlap of genera and variation in the relative abundance resulted in a statistically-significant difference in their composition (PERMANOVA, p = 0.028 & F = 2.059) but not in the plant taxon dispersion (PERMADISP, p = 0.77 & F = 0.089) **(Figure 4b)**. This suggests that the variability of the composition of individual samples within each hive are similar but there are overall significant differences among the two hives.

**Figure 4:**
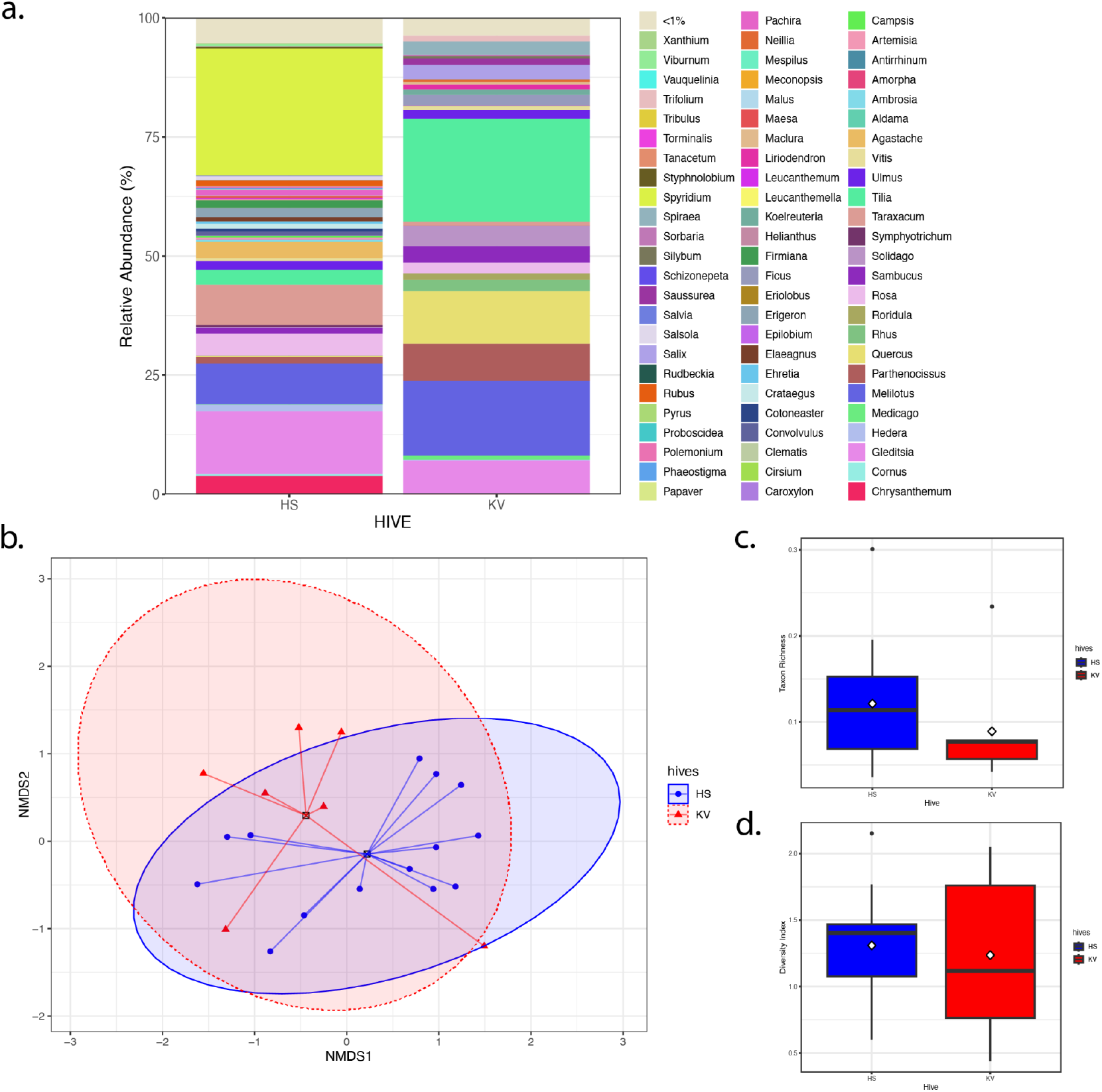
Comparison of the Total HS and KV Hive’s Pollen Composition. (a) Stacked bar charts illustrating the taxonomic composition (at the genus level) of pollen samples collected from the two hives. The RRA of each taxon is determined by the pooling of all samples from their respective hives (HS, n = 14 & KV, n = 7). Taxa accounting for <1% of reads in each hive are grouped as “ <1%”. (b) Composition of plant genera in the KV hive (red, n = 7) and HS hive (blue, n = 14). This non-metric multidimensional scaling (NMDS) plot is based on Hellinger Dissimilarity Index Data and ellipses show the 95% confidence level for the two hives. (c) Boxplots of the taxa richness for the HS hive (blue, n = 14) and KV hive (red, n = 7). (d) Boxplots of the Shannon diversity index for the HS hive (blue, n = 14) and KV hive (red, n = 7).

There was not a statistical difference in the taxa richness between the two hives (Mann-Whitney U-Test, p = 0.2545, W = 65). HS samples’ richness ranged from 0.036 to 0.30 and the KV samples ranged from 0.042 to 0.23 **(Figure 4c)**. The Shannon diversity index also did not differ significantly (Mann-Whitney U-Test, p = 0.7433, W = 54) ranging from 0.60 to 2.15 in the HS hive and from 0.44 to 2.05 in the KV hive **(Figure 4d)**.

Samples taken from both hives on June 6th, June 20th, June 27th, July 25th, and September 12th allow us to examine how the two hives differ within the course of month (June) as well as throughout the entire season (June - Early, July - Mid, and September - Late) **(Figure 5a-b)**. While the difference in taxa richness of the samples ranged from 0.0058 (06/06) to 0.24 (06/20), on a given day, the diversity indexes of the two hives followed very similar trends (largest difference in richness being 0.73 on 9/12). Both hives had maximum Shannon diversity on June 20th (HS = 2.15 & KV = 2.05) and minimum Shannon diversity on July 25th (HS = 0.695 & KV = 0.44) **(Figure 5c)**.

**Figure 5:**
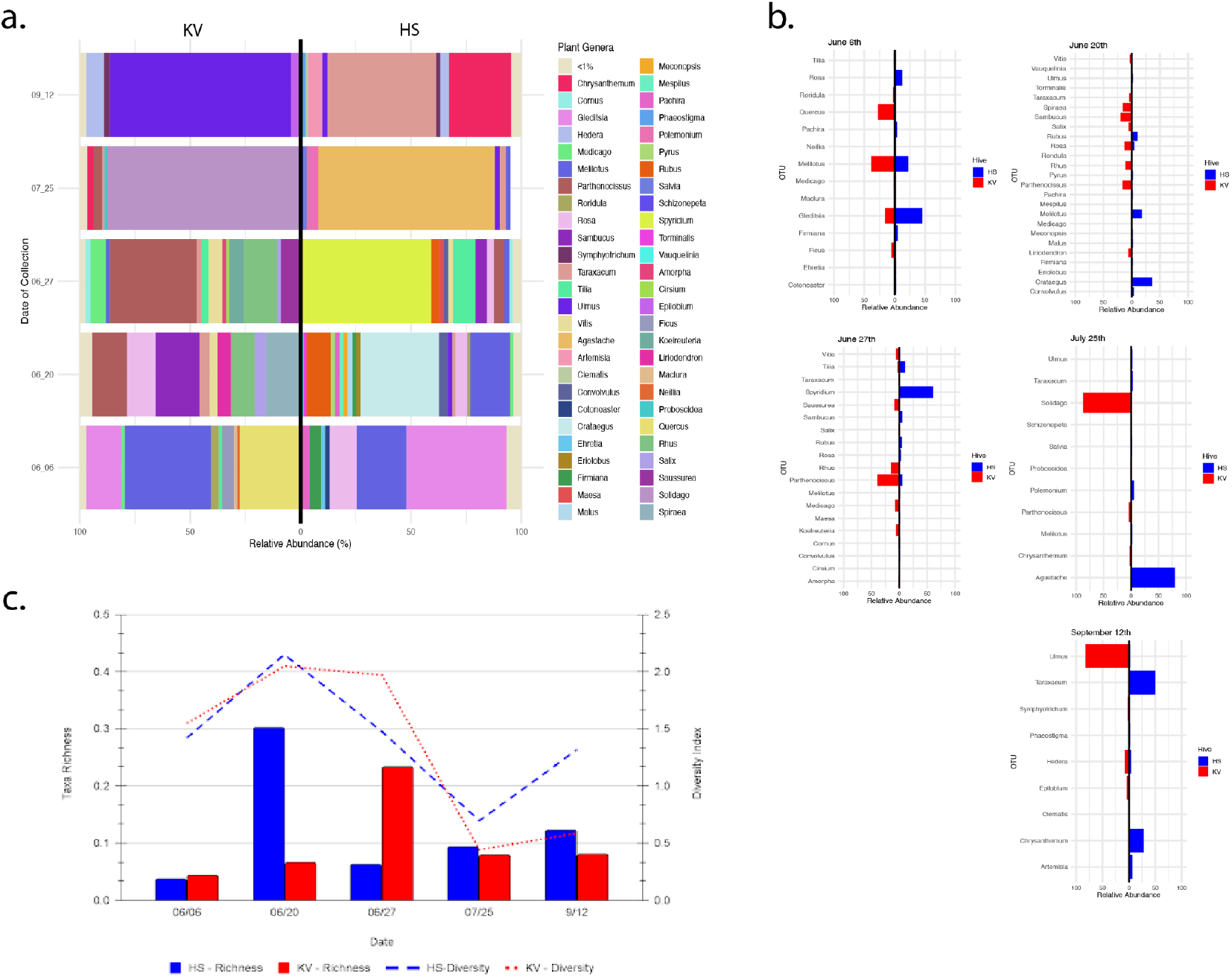
Comparison of HS and KV Pollen on 5 Sampling Days. (a) Mirror, stacked, bar graph shows the taxonomic composition (at the genus level) of the HS hive (left) and KV hive (right) on 5 separate days. The relative abundance is based on the percent composition of each taxa. (b) Mirrored bar graph shows the relative abundance of taxa present in the HS hive (blue) and/or the KV hive (red). (c) Mixed graph showing the taxa richness (bars) and Shannon diversity indexes (line) of the HS (blue) and KV (red) hives on five separate days throughout the season.

Despite the similarity in key diversity index trends, the composition of samples taken on the same day display large levels of taxonomic diversity. The percentage of taxa that were identified in both hives over the total number of taxa identified ranges from 8% (06/20 samples) to 33.3% (09/12 samples). The first four pairwise comparisons between samples taken on the same day did not meet thresholds for significance (p > 0.5) with Spearman Rho ranging between -0.113 and 0.247 **(Table S5**). The only sampling day that resulted in a significant correlation in plant composition was September 9th with a Spearman Rho of 0.454 (p = 0.001) and a shared taxa percentage of 33.3%.

## Discussion

This study aims to apply metabarcoding and next generation sequencing techniques to identify plant taxa present in corbicular pollen samples. The collection and subsequent identification of these plant taxa fulfills two major aims: 1) determine trends in foraging behavior from one hive across the foraging season 2.) investigate the similarity between the foraging of two closely related hives. Due to Apis mellifera’s crucial role in agriculture and local ecosystems, it is becoming more important than ever before to understand the factors that contribute to pollen collection due to the important nutritional role that it plays in hive health. The use of NGS sequencing technologies such as ONT provides relatively quick and accurate descriptions of in-depth compositional analysis and allows for the undertaking of more expansive and longitudinal studies in the future.

A total of seventy-four taxa (at the genus level) were identified across a total of twenty-one samples collected in this study. Many of these taxa were also identified in other papers studying *Apis mellifera* foraging behaviors across the world. For instance, *Quercus* and *Trifolium* were identified using both microscopy on pollen collected from Texas and California and metabarcoding techniques on bee bread and honey from hives located in Finland (Lau et al., 2018; Leponiemi et al., 2023). Other prominent plant genera are identifiable in the Salt Lake City area and are native and/or ornamental flowering plants, shrubs, and trees (https://extension.usu.edu/ecorestore/ & https://facilities.utah.edu/tree-tour/). The prevalence of the identified taxa in these other studies and surveys suggests that this method of taxonomic identification is a valid measure of plant taxa presence in pollen samples.

### Pollen Foraging Varies throughout the Course of a Foraging Season

Across the foraging season, 59 distinct taxonomic units (genus level) were identified in HS hive samples. The majority of these 59 taxa encompassed small percentages of the samples in which they were identified with only 25 taxa identified at over 5% relative read abundances in any of the fourteen samples. In addition, only 5 taxa encompassed over 50% of the reads in a single sample. This suggests that although a median of 8 distinct taxa were identified per sample, on average a single taxonomic unit encompassed around 54.68% of the sample (minimum of 28.94% and maximum of 80.23%). This result supports both A. mellifera’s generalist foraging behavior while simultaneously suggesting a degree of pollen preference.

An important consideration is the relationship between floral availability and taxonomic diversity and richness. Throughout the season, the taxa richness fluctuated from 0.036 to 0.301. This fluctuation occurred over the course of a two-week time period. Following this peak, richness experienced a hard decline the following week with a gradual increase until early August when the trend sloped downward for the remainder of the season. The Shannon diversity index which accounts for both richness and evenness followed a similar trend but started relatively high hitting its peak on the same collection day as richness (2.15 on 06/20) but its minimum on the last day of collection (0.601 on 10/17). The similarity in these two primary measures of diversity indicates that late June and early August both experienced an increase in the number of foraging targets as well as a more even distribution of the proportion of pollen collected from each source. This increased diversity in June and subsequent decrease aligns with pollen’s genus richness observed across 12 apiaries in Pennsylvania (Sponsler et al., 2020).

By breaking the observed foraging season into three temporal groups, it is evident that the number of taxa pollen collected decreases throughout the summer. Across the five-week period at the beginning of the collection season (labeled Early Season), 33 taxa were identified. During the next collection season which spans five weeks in late July and early August (labeled as Mid Season) the number of taxa identified decreases to 26. Finally, across the last four samples spanning the last 6 weeks of the collection period season, only 13 taxa were identified. Despite this substantial decrease in the number of plant taxa present in September and October, 69.2% (9 of the 13 taxonomic groups) were present in one or both of the other seasons. This is vastly different compared to the early season which only shared 15.2% (5 of 33 taxa) with the other seasons. This difference in the identification of taxa present suggests that the foraging behavior of A. mellifera in early summer is distinct compared to later summer foraging which is relatively similar.

With the diversity of plant taxa identified throughout the season (59) being so large, it is somewhat unexpected that not a single plant taxa was identified in all fourteen samples. Taraxacum (a genus of the family Asteraceae containing many species of dandelions) was the most prevalent plant genera identified being one of three taxa present in all three temporal seasons. Even then it was not identified in relative abundances over 1% in four of the five Early Season samples. This demonstrates the high level of variability in pollen collection across time but the relatively low number of taxa present in individual samples suggest a lack of variability in foraging on a given day.

### Hives within the Same Foraging Range have Distinct Pollen Foraging Preferences

Comparisons of the two hives’ pollen indicate significant differences in plant composition. While there was no significant difference observed in the dispersion, average taxonomic richness, or diversity indexes, there was a significant difference in their respective plant compositions. Across the 59 taxa identified in the fourteen HS samples and the 33 taxa identified in the seven KV samples, eighteen of these taxa were identified in both hives’ pollen samples. Despite this 54.5% shared taxa for KV and 30.5% shared taxa for HS, the relative abundance of this taxa tells a much different story. While many key contributors of one hive are present in the other hive, the relative abundance of these genera varies drastically. This variability includes Gleditsia and Tilia (the most prevalent shared taxa in the HS hive and KV hive, respectively) which encompass 13.14% and 3.13% of the HS hives’ total composition respectively but encompass only 6.88% and 21.57% of the KV hives’ composition.

This difference between these two hives can also be seen on a daily basis. Five out of the nineteen total collection days resulted in sufficient pollen being collected from both hives. Among these five collection days, only one day (09/12) had significantly correlated compositions of the two hives. The remaining days showed no correlation. Given that these hives are well within A. mellifera’s average foraging range, several other aspects could be accumulating to result in this drastic difference. These alterations in pollen foraging may be attributed to brood size, queen presence, urbanization, homogeneity of plant distribution, and distance to resources (Davey et al., 2024; Free, 1967). While this study on its own is unable to draw any conclusions regarding the causation behind these differences in pollen foraging, it does demonstrate the need for further longitudinal studies regarding pollen foraging behavior using metabarcoding techniques.

## Notes

### Competing Interest Statement

The authors have declared no competing interest.

