## Supplementary material for "Investigating Honey Bee Pollen Foraging Patterns Across a Season Using DNA Metabarcoding": Figure S1

**Figure S1: Hierarchical Clustering of the HS samples**. A Euclidean distance–based dendrogram that summarizes the differentiation among pollen samples taken from the HS hives separated into Early (red), Mid (blue), and Late (green) Seasons based on the date of collection.

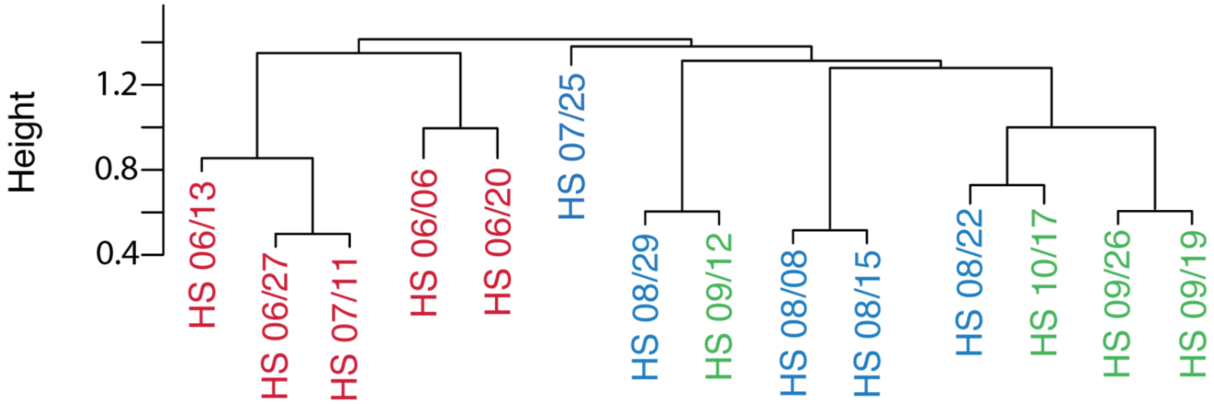
