## Supplementary material for "Investigating Honey Bee Pollen Foraging Patterns Across a Season Using DNA Metabarcoding": Table S1

**Table S1: Multiplexing, Metabarcoding Primer Sequences.** The lowercase nucleotides represent the index while the uppercase nucleotides represent the primer sequence.

| Primers | Sequence |
|---|---|
| trnL_C_F1 | agttatcgttcgaggtgttcagtgCGAAATCGGTAGACGCTACG |
| trnL_C_F2 | atggtcggctcgatgtcatgaataCGAAATCGGTAGACGCTACG |
| trnL_C_F3 | atgtgccgggattgaaatacgtccCGAAATCGGTAGACGCTACG |
| trnL_C_F4 | cactacctcgttcaagcttacatcCGAAATCGGTAGACGCTACG |
| trnL_D_R1 | ggtaaagggctatcttgtgttgcgGGGGATAGAGGGACTTGAAC |
| trnL_D_R2 | tcgctctgttatctaaaggccgtaGGGGATAGAGGGACTTGAAC |
| trnL_D_R3 | tggtagattaactcgaacatctccGGGGATAGAGGGACTTGAAC |
| trnL_D_R4 | tggtattgagcacctaagactggaGGGGATAGAGGGACTTGAAC |
