## Supplementary material for "Investigating Honey Bee Pollen Foraging Patterns Across a Season Using DNA Metabarcoding": Table S2

**Table S2: HS Hive RRA Values.** RRA values above the "<1" cutoff are highlighted in gray while genera present in less that the cutoff are marked as "<1%" and genera not identified in the corresponding sample (HS - n/n/2023) are marked as "N/A".

| | **HS - 06/06** | **HS - 06/13** | **HS - 06/20** | **HS - 06/27** | **HS - 07/11** | **HS - 07/25** | **HS - 08/08** | **HS - 08/15** | **HS - 08/22** | **HS - 08/29** | **HS - 09/12** | **HS - 09/19** | **HS - 09/26** | **HS - 10/17** |
|---|---|---|---|---|---|---|---|---|---|---|---|---|---|---|
| **Agastache** | N/A | <1% | N/A | N/A | <1% | 80.23 | 1.32 | <1% | <1% | N/A | <1% | <1% | 2.23 | N/A |
| **Aldama** | N/A | N/A | N/A | N/A | N/A | N/A | <1% | 1.36 | <1% | N/A | <1% | N/A | N/A | N/A |
| **Ambrosia** | N/A | <1% | N/A | N/A | N/A | N/A | <1% | 5.98 | 8.58 | <1% | <1% | <1% | <1% | N/A |
| **Antirrhinum** | N/A | N/A | N/A | N/A | N/A | N/A | <1% | 2.96 | N/A | N/A | N/A | N/A | <1% | N/A |
| **Artemisia** | N/A | N/A | N/A | N/A | <1% | N/A | <1% | <1% | <1% | 5.82 | 6.51 | <1% | <1% | <1% |
| **Campsis** | N/A | N/A | <1% | N/A | <1% | N/A | <1% | <1% | 7.25 | 1.62 | <1% | <1% | <1% | N/A |
| **Caroxylon** | N/A | N/A | N/A | N/A | N/A | N/A | 1.04 | <1% | <1% | <1% | <1% | N/A | N/A | N/A |
| **Chrysanthemum** | N/A | <1% | N/A | <1% | 1.28 | <1% | 4.10 | <1% | <1% | 70.08 | 28.40 | <1% | <1% | <1% |
| **Clematis** | N/A | N/A | N/A | N/A | N/A | <1% | N/A | N/A | N/A | <1% | 1.19 | N/A | N/A | N/A |
| **Convolvulus** | <1% | <1% | 3.87 | 1.81 | 2.64 | <1% | N/A | N/A | <1% | <1% | N/A | <1% | <1% | <1% |
| **Cornus** | <1% | <1% | <1% | 1.54 | <1% | N/A | N/A | N/A | <1% | N/A | N/A | N/A | N/A | <1% |
| **Cotoneaster** | 2.01 | <1% | <1% | <1% | <1% | <1% | N/A | N/A | N/A | N/A | N/A | N/A | N/A | N/A |
| **Crataegus** | <1% | 2.56 | 35.89 | <1% | <1% | <1% | N/A | <1% | N/A | N/A | <1% | <1% | N/A | <1% |
| **Ehretia** | 1.83 | N/A | N/A | <1% | <1% | N/A | N/A | N/A | N/A | N/A | N/A | N/A | N/A | N/A |
| **Elaeagnus** | <1% | 28.94 | <1% | <1% | <1% | <1% | N/A | <1% | <1% | N/A | <1% | <1% | <1% | <1% |
| **Erigeron** | N/A | <1% | <1% | N/A | N/A | <1% | 40.77 | 52.04 | <1% | <1% | <1% | <1% | <1% | N/A |
| **Eriolobus** | <1% | <1% | 1.68 | N/A | N/A | <1% | N/A | N/A | N/A | N/A | N/A | <1% | N/A | <1% |
| **Firmiana** | 5.18 | <1% | 1.92 | <1% | <1% | N/A | N/A | <1% | N/A | N/A | <1% | <1% | <1% | <1% |
| **Gleditsia** | 45.31 | 4.19 | <1% | <1% | 1.24 | N/A | N/A | N/A | N/A | N/A | N/A | N/A | <1% | <1% |
| **Hedera** | N/A | <1% | <1% | N/A | N/A | <1% | <1% | <1% | <1% | <1% | 3.97 | 7.41 | 40.30 | 2.07 |
| **Helianthus** | N/A | N/A | N/A | N/A | N/A | N/A | <1% | <1% | 2.27 | <1% | <1% | <1% | N/A | N/A |
| **Leucanthemella** | N/A | N/A | N/A | N/A | <1% | N/A | <1% | N/A | N/A | 2.75 | <1% | N/A | <1% | N/A |
| **Leucanthemum** | N/A | N/A | N/A | <1% | 1.97 | N/A | N/A | N/A | N/A | N/A | N/A | N/A | N/A | N/A |

| | | | | | | | | | | | | | | |
|---|---|---|---|---|---|---|---|---|---|---|---|---|---|---|
| **Maesa** | <1% | <1% | N/A | 1.88 | <1% | N/A | N/A | <1% | N/A | N/A | N/A | N/A | N/A | N/A |
| **Malus** | <1% | <1% | 2.22 | N/A | N/A | <1% | N/A | N/A | N/A | N/A | N/A | N/A | <1% | N/A |
| **Meconopsis** | N/A | N/A | 1.98 | <1% | N/A | N/A | N/A | N/A | N/A | N/A | N/A | <1% | N/A | <1% |
| **Medicago** | <1% | 1.72 | 1.43 | <1% | <1% | <1% | <1% | <1% | <1% | <1% | <1% | <1% | <1% | <1% |
| **Melilotus** | 22.45 | 28.68 | 18.16 | 2.22 | 1.50 | 2.00 | <1% | <1% | <1% | <1% | <1% | <1% | <1% | <1% |
| **Mespilus** | <1% | <1% | 1.65 | N/A | N/A | <1% | N/A | <1% | N/A | N/A | N/A | N/A | <1% | <1% |
| **Pachira** | 4.10 | <1% | 2.03 | <1% | <1% | N/A | N/A | <1% | <1% | N/A | <1% | <1% | N/A | <1% |
| **Papaver** | N/A | 2.08 | N/A | N/A | N/A | N/A | N/A | N/A | N/A | N/A | N/A | N/A | <1% | N/A |
| **Parthenocissus** | <1% | <1% | <1% | 4.85 | 2.09 | N/A | <1% | <1% | <1% | <1% | N/A | N/A | <1% | <1% |
| **Phaeostigma** | N/A | <1% | N/A | <1% | <1% | N/A | <1% | <1% | <1% | 4.45 | 2.23 | <1% | <1% | <1% |
| **Polemonium** | N/A | N/A | <1% | N/A | N/A | 5.14 | 6.83 | 1.60 | <1% | <1% | N/A | N/A | N/A | <1% |
| **Pyrus** | <1% | <1% | 2.14 | N/A | N/A | <1% | N/A | N/A | N/A | N/A | N/A | <1% | N/A | <1% |
| **Quercus** | <1% | N/A | N/A | <1% | <1% | <1% | <1% | <1% | <1% | <1% | <1% | <1% | 3.62 | <1% |
| **Rhus** | <1% | 1.46 | <1% | <1% | <1% | N/A | N/A | N/A | <1% | N/A | N/A | N/A | <1% | <1% |
| **Roridula** | <1% | 1.63 | 1.38 | <1% | <1% | <1% | N/A | N/A | N/A | <1% | N/A | <1% | <1% | N/A |
| **Rosa** | 12.37 | <1% | 5.28 | 2.99 | <1% | <1% | N/A | <1% | <1% | <1% | <1% | <1% | N/A | 11.38 |
| **Rubus** | <1% | <1% | 10.91 | 3.86 | <1% | N/A | N/A | <1% | <1% | N/A | <1% | <1% | N/A | <1% |
| **Rudbeckia** | N/A | N/A | N/A | N/A | <1% | <1% | 1.19 | 1.70 | N/A | <1% | <1% | N/A | N/A | N/A |
| **Salsola** | N/A | <1% | N/A | N/A | N/A | N/A | 21.78 | 10.88 | 2.96 | <1% | <1% | N/A | <1% | <1% |
| **Salvia** | N/A | N/A | N/A | N/A | N/A | 1.69 | N/A | N/A | N/A | N/A | N/A | N/A | N/A | N/A |
| **Sambucus** | <1% | N/A | N/A | 5.29 | <1% | N/A | N/A | <1% | <1% | N/A | <1% | <1% | N/A | N/A |
| **Schizonepeta** | N/A | N/A | N/A | N/A | N/A | 1.10 | N/A | N/A | N/A | N/A | <1% | N/A | <1% | N/A |
| **Solidago** | N/A | <1% | <1% | N/A | N/A | <1% | 2.88 | 1.97 | <1% | 2.15 | <1% | <1% | <1% | <1% |
| **Spyridium** | <1% | 10.17 | <1% | 59.26 | 78.98 | N/A | N/A | <1% | N/A | <1% | <1% | <1% | <1% | <1% |
| **Styphnolobium** | <1% | <1% | N/A | N/A | N/A | <1% | 5.94 | <1% | 7.38 | <1% | N/A | N/A | <1% | N/A |
| **Symphyotrichum** | N/A | N/A | <1% | N/A | N/A | <1% | <1% | <1% | 1.62 | <1% | 1.74 | 1.09 | 5.29 | 3.08 |
| **Tanacetum** | N/A | <1% | N/A | <1% | <1% | N/A | 2.11 | <1% | <1% | 1.00 | <1% | <1% | <1% | N/A |

| | | | | | | | | | | | | | | |
|---|---|---|---|---|---|---|---|---|---|---|---|---|---|---|
| **Taraxacum** | <1% | <1% | 1.41 | <1% | <1% | 2.79 | 9.04 | 16.96 | 58.35 | 7.18 | 49.69 | 34.79 | 34.74 | 79.82 |
| **Tilia** | <1% | 14.88 | <1% | 10.19 | 2.14 | N/A | N/A | <1% | <1% | N/A | <1% | <1% | <1% | <1% |
| **Torminalis** | <1% | <1% | 1.33 | N/A | N/A | <1% | N/A | N/A | N/A | N/A | N/A | <1% | N/A | <1% |
| **Tribulus** | N/A | <1% | <1% | <1% | N/A | <1% | N/A | N/A | N/A | N/A | <1% | <1% | 2.04 | N/A |
| **Ulmus** | <1% | <1% | 2.17 | <1% | N/A | 2.17 | <1% | <1% | <1% | <1% | 2.18 | 45.83 | 8.19 | <1% |
| **Vauquelinia** | <1% | <1% | 1.35 | <1% | <1% | <1% | N/A | <1% | N/A | N/A | <1% | <1% | <1% | <1% |
| **Viburnum** | <1% | <1% | <1% | <1% | <1% | <1% | N/A | <1% | 2.00 | <1% | <1% | 8.47 | <1% | <1% |
| **Vitis** | <1% | <1% | <1% | 2.26 | <1% | N/A | N/A | N/A | <1% | <1% | N/A | N/A | <1% | <1% |
| **Xanthium** | <1% | N/A | <1% | <1% | <1% | <1% | <1% | <1% | 5.02 | <1% | <1% | N/A | <1% | <1% |
