## Supplementary material for "Investigating Honey Bee Pollen Foraging Patterns Across a Season Using DNA Metabarcoding": Table S3

**Table S3: KV Hive RRA Values.** RRA values above the "<1" cutoff are highlighted in gray while genera present in less that the cutoff are marked as "<1%" and genera not identified in the corresponding sample (KV - n/n/2023) are marked as "N/A".

| | KV - 06/06 | KV - 06/20 | KV - 06/27 | KV - 07/06 | KV - 07/18 | KV - 07/25 | KV - 09/12 |
|---|---|---|---|---|---|---|---|
| **Amorpha** | N/A | N/A | 1.93 | <1% | <1% | N/A | N/A |
| **Chrysanthemum** | <1% | <1% | <1% | <1% | 1.18 | 2.90 | <1% |
| **Cirsium** | N/A | N/A | 1.20 | <1% | N/A | N/A | N/A |
| **Cornus** | <1% | N/A | 2.18 | N/A | N/A | <1% | N/A |
| **Epilobium** | N/A | N/A | N/A | N/A | N/A | N/A | 4.26 |
| **Ficus** | 5.47 | <1% | N/A | <1% | N/A | N/A | N/A |
| **Gleditsia** | 16.00 | <1% | <1% | 1.50 | <1% | N/A | N/A |
| **Hedera** | N/A | N/A | <1% | N/A | <1% | N/A | 8.22 |
| **Koelreuteria** | <1% | <1% | 6.60 | 3.16 | <1% | <1% | N/A |
| **Liriodendron** | N/A | 5.96 | N/A | <1% | <1% | N/A | N/A |
| **Maclura** | 1.43 | <1% | N/A | <1% | N/A | N/A | N/A |
| **Medicago** | 1.62 | <1% | 7.23 | <1% | <1% | <1% | <1% |
| **Melilotus** | 39.19 | <1% | 1.64 | <1% | <1% | <1% | <1% |
| **Neillia** | 1.17 | <1% | N/A | <1% | N/A | N/A | N/A |
| **Parthenocissus** | <1% | 15.92 | 39.36 | 5.03 | 60.35 | 3.91 | <1% |
| **Proboscidea** | <1% | N/A | <1% | <1% | N/A | 1.52 | N/A |
| **Quercus** | 27.56 | <1% | <1% | <1% | N/A | N/A | N/A |
| **Rhus** | N/A | 10.95 | 15.25 | <1% | N/A | <1% | N/A |
| **Roridula** | 3.41 | <1% | <1% | <1% | N/A | <1% | <1% |
| **Rosa** | <1% | 12.99 | <1% | <1% | <1% | <1% | <1% |
| **Salix** | <1% | 5.06 | 1.55 | 6.87 | N/A | N/A | N/A |
| **Sambucus** | <1% | 19.88 | N/A | <1% | N/A | N/A | N/A |
| **Saussurea** | <1% | <1% | 9.00 | 3.71 | <1% | <1% | N/A |
| **Silybum** | <1% | <1% | <1% | 1.52 | <1% | N/A | N/A |
| **Solidago** | N/A | N/A | <1% | N/A | 28.49 | 87.61 | N/A |
| **Sorbaria** | <1% | N/A | N/A | 1.12 | N/A | N/A | N/A |
| **Spiraea** | <1% | 15.63 | N/A | <1% | N/A | N/A | N/A |
| **Symphyotrichum** | N/A | N/A | <1% | N/A | <1% | <1% | 2.65 |
| **Taraxacum** | <1% | 4.59 | 1.89 | <1% | <1% | 1.03 | <1% |
| **Tilia** | 1.51 | <1% | 3.47 | 68.94 | <1% | <1% | <1% |
| **Trifolium** | <1% | N/A | <1% | 3.24 | 6.16 | <1% | N/A |
| **Ulmus** | <1% | N/A | <1% | N/A | <1% | <1% | 82.45 |
| **Vitis** | <1% | 3.59 | 6.19 | <1% | 1.27 | <1% | N/A |
