## Supplementary material for "Investigating Honey Bee Pollen Foraging Patterns Across a Season Using DNA Metabarcoding": Table S4

**Table S4: Shared Taxa between the HS and KV Hives.** Plant genera present in only the HS hive samples (left), only the KV hive samples (right), or identified in samples from both hives (middle). To be considered, taxa must be identified at an RRA of ≥1% in 1 or more samples from the corresponding hive.

| **Only HS** | **HS AND KV** | **Only KV** |
|---|---|---|
| Agastache | Chrysanthemum | Amorpha |
| Aldama | Cornus | Cirsium |
| Ambrosia | Gleditsia | Epilobium |
| Antirrhinum | Hedera | Ficus |
| Artemisia | Medicago | Koelreuteria |
| Campsis | Melilotus | Liriodendron |
| Caroxylon | Parthenocissus | Maclura |
| Clematis | Quercus | Neillia |
| Convolvulus | Rhus | Proboscidea |
| Cotoneaster | Roridula | Salix |
| Crataegus | Rosa | Saussurea |
| Ehretia | Sambucus | Silybum |
| Elaeagnus | Solidago | Sorbaria |
| Erigeron | Symphyotrichum | Spiraea |
| Eriolobus | Taraxacum | Trifolium |
| Firmiana | Tilia | |
| Helianthus | Ulmus | |
| Leucanthemella | Vitis | |
| Leucanthemum | | |
| Maesa | | |
| Malus | | |
| Meconopsis | | |
| Mespilus | | |
| Pachira | | |
| Papaver | | |
| Phaeostigma | | |
| Polemonium | | |
| Pyrus | | |
| Rubus | | |
| Rudbeckia | | |
| Salsola | | |
| Salvia | | |
| Schizonepeta | | |
| Spyridium | | |

| |
|---|
| Styphnolobium |
| Tanacetum |
| Torminalis |
| Tribulus |
| Vauquelinia |
| Viburnum |
| Xanthium |
