## Supplementary material for "Investigating Honey Bee Pollen Foraging Patterns Across a Season Using DNA Metabarcoding": Table S5

**Table S5: Spearman Correlation Values.** These values are based on correlation analysis between HS and KV samples collected on the same day. The shared taxa percentage is calculated using the number of total taxa identified across the two samples and the number of these taxa identified at an RRA ≥1%.

| Date | Spearman Rho | Spearman P-Value | Shared Taxa (%) |
|---|---|---|---|
| 6/6/2023 | 0.183 | 0.189 | 14.3% |
| 6/20/2023 | -0.113 | 0.421 | 8% |
| 6/27/2023 | 0.247 | 0.074 | 26.3% |
| 7/25/2023 | 0.057 | 0.687 | 9.1% |
| 9/12/2023 | 0.454 | 0.001 | 33.3% |
